# Highly efficient CRISPR/Cas9-mediated tissue specific mutagenesis in *Drosophila*

**DOI:** 10.1101/268482

**Authors:** Amy R. Poe, Bei Wang, Maria L. Sapar, Hui Ji, Kailyn Li, Tireniolu Onabajo, Rushaniya Fazliyeva, Mary Gibbs, Yue Qiu, Yuzhao Hu, Chun Han

## Abstract

Tissue-specific loss-of-function (LOF) analysis is an essential approach for characterizing gene function. Here we describe an efficient <u>CRISPR</u>-mediated <u>t</u>issue-<u>r</u>estricted <u>m</u>utagenesis (CRISPR-TRiM) method for ablating gene function in *Drosophila*. This binary system consists of a tissue-specific Cas9 and a ubiquitously expressed multi-guide RNA (gRNA) transgene. To facilitate the construction of these components, we developed convenient tools for generating and evaluating enhancer-driven Cas9 lines, identified a multi-gRNA design that is highly efficient in mutagenizing somatic cells, and established an assay for testing the efficiency of multi-gRNAs in creating double-stranded breaks. We found that excision of genomic DNA induced by two gRNAs is infrequent in somatic cells, while indels more reliably cause tissue-specific LOF. Furthermore, we show that enhancer-driven Cas9 is less cytotoxic yet results in more complete gene removal than Gal4-driven Cas9 in larval neurons. Finally, we demonstrate that CRISPR-TRiM efficiently unmasks redundant gene functions in neuronal morphogenesis. Importantly, two Cas9 transgenes that turn on with different timings in the neuronal lineage revealed the extent to which gene products persist in cells after tissue-specific gene knockout. These CRISRPR tools can be applied to analyze tissue-specific gene function in many biological processes.

## INTRODUCTION

Tissue-specific loss-of-function (LOF) analysis is instrumental for elucidating the developmental roles of essential genes, determining cell autonomy, and dissecting cell-cell interactions. Conventional methods for studying tissue-specific gene function in *Drosophila*, such as mosaic analysis with a repressible cell marker (MARCM)^1^ and tissue-specific RNA interference (RNAi)^2,3^,are powerful approaches for genetic screens and LOF analysis. However, these techniques present several disadvantages. RNAi is prone to off-target effects^4^ and gene knockdown is rarely complete^2^ because this technique only targets mRNAs for degradation or translational suppression. MARCM produces more reliable LOF of genes of interest, but the process can be labor intensive and requires multiple components to be combined in the same animal.

The CRISPR/Cas9 system^5^ has the potential to surpass the current methods of tissue-specific LOF in *Drosophila* due to its simplicity and efficiency in creating gene disruption^6–11^. In this system, Cas9 endonuclease cleaves double-stranded genomic DNA at a site determined by the protospacer sequence (or targeting sequence) of a chimeric guide RNA (gRNA)5. Cas9-mediated double-stranded breaks (DSBs) at the targeting site are then repaired by the host through either nonhomologous end joining (NHEJ) or homology-directed repair (HDR)^12^. Imprecise repair through NHEJ can result in small insertions or deletions (indels) at a single target site^6^ or deletions of DNA fragments between two target sites^9,10^. CRISPR/Cas9 has been successfully used in *Drosophila* and other organisms to create heritable mutations^6–8^, to edit genomic sequences precisely ^13,14^, and to control gene expression^15,16^.

Conditional mutagenesis of genes has been achieved in *Drosophila* by combining the CRISPR/Cas9 system with the Gal4/UAS system^17–19^. In this approach, tissue-specific Gal4 drives *UAS-Cas9* expression, while gRNAs are expressed either from ubiquitous promoters^17,18^ or by UAS^19^. Transgenic constructs expressing multiple gRNAs increase mutagenesis efficiency and allow simultaneous mutagenesis of more than one gene^17–19^. Despite these initial successes, Gal4-driven Cas9 and transgenic gRNAs have not been widely used to study tissue-specific gene function due to uncertainties and limitations associated with this method. For example, gRNAs can vary greatly in their mutagenic efficiency, and it is difficult to know whether a transgenic gRNA reliably causes mutations in the tissue of interest. These concerns worsen when a multiplex gRNA construct is used to knock out two or more genes simultaneously. Gal4-driven Cas9 has several additional potential drawbacks that could limit its applications in developmental studies. First, the intermediate Gal4 expression step can delay Cas9 expression, making it difficult to study early gene functions in specific tissues. Second, the Gal4/UAS system often results in excessive levels of Cas9 expression which can be toxic^20^. Finally, using Gal4-driven Cas9 makes the Gal4/UAS system unavailable for other genetic manipulations in the same animal. Thus, a simpler and more robust method of tissue-specific mutagenesis is needed to take full advantage of the CRISPR/Cas9 system.

One way to improve conditional mutagenesis is optimization of transgenic gRNAs. The mutagenic efficiency of a gRNA is affected by both the gRNA target sequence and the transgenic vector design. Previous studies in *Drosophila* exploring choices of the gRNA promoter, the length and sequence composition of the target sequence, and methods of producing multiple gRNAs from a single construct have identified several parameters for making efficient gRNAs^17–19,21^. However, the goal of most of these strategies was to increase the frequency of heritable mutations, leaving room for optimization of transgenic gRNA design for mutagenesis in somatic cells. In addition, specific modifications of the gRNA scaffold improve Cas9 targeting to DNA in human cells^22^, but these modifications have not been tested to date in *Drosophila*. Thus, there is a compelling need for optimized transgenic gRNAs coupled with tissue-specific control of Cas9 efficacy.

Here, we have developed a new CRISPR/Cas9 toolkit that achieves highly efficient knockout of one or multiple *Drosophila* genes in a tissue-specific manner. Our method of <u>CRISPR</u>-mediated tissue-<u>r</u>estricted <u>m</u>utagenesis (CRISPR-TRiM) (Fig. 1a) combines a transgenic Cas9 driven by a tissue specific enhancer with a transgenic construct that ubiquitously expresses multiple gRNAs. By targeting every gene of interest with two gRNAs, this system mutates all target genes tissue-specifically through indel formation or large DNA deletions. To build the most efficient reagents, we have generated convenient tools for making and evaluating enhancer-driven Cas9 transgenes, identified a multi-gRNA design that is superior to previous options, and established an *in vivo* assay for testing gRNA efficiency in causing DSBs. We investigated how the frequency of DNA deletion in individual somatic cells is impacted by the distance between two target sites and we further found that enhancer-driven Cas9 is more effective in causing LOF and less cytotoxic than Gal4-driven Cas9.Using genes in the SNARE (soluble *N*-ethylmalemide–sensitive factor attachment protein receptor) pathway as examples, we demonstrate here that CRISPR-TRiM can efficiently knock out multiple redundant genes in neurons. Our results also underscore the importance of mutagenesis timing for uncovering tissue-specific gene functions: Post-mitotic knockout of neuronal type-specific genes, such as the receptor protein tyrosine phosphatase *Ptp69D*, is sufficient and effective for removing gene functions; while housekeeping genes, such as those encoding NSF and SNAP proteins, require mutagenesis earlier in the cell lineage to unmask their LOF phenotypes.

**Fig. 1.**
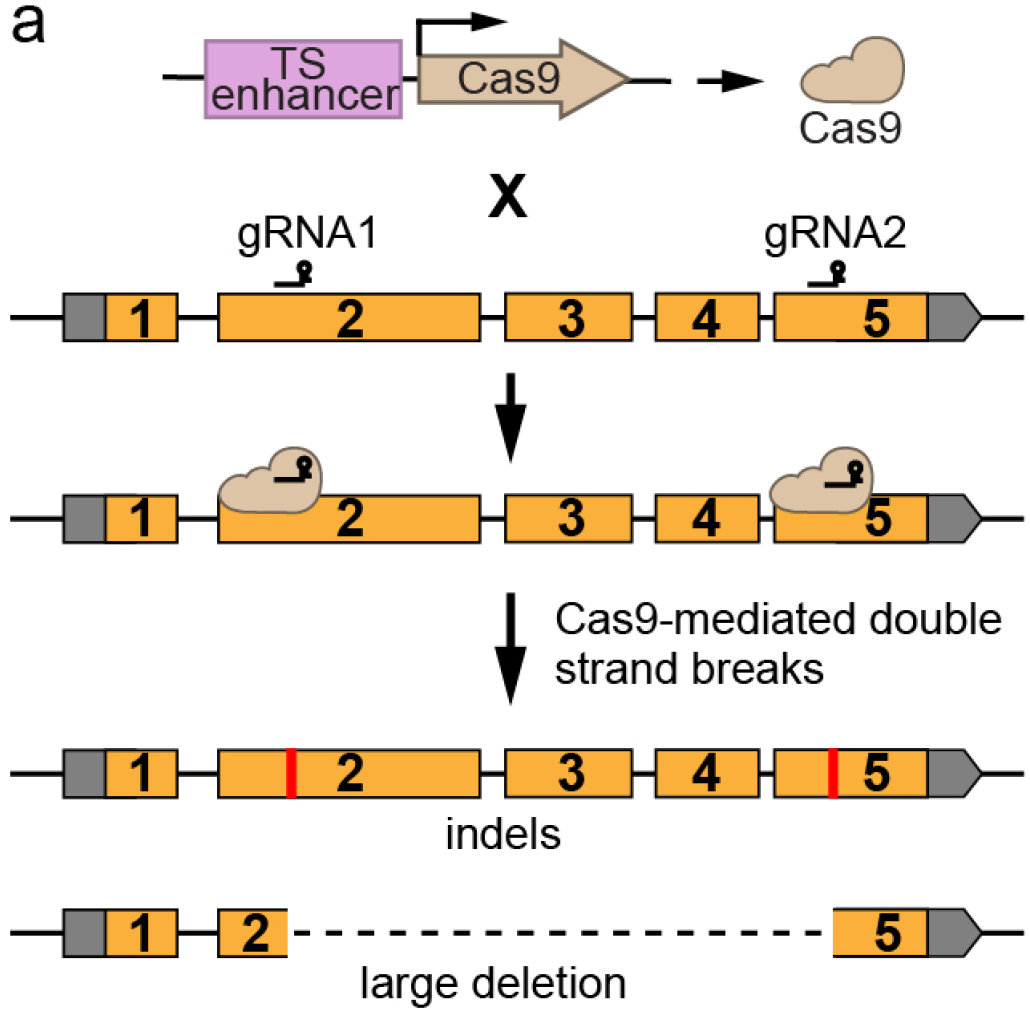
A diagram of CRISPR-TRiM. A tissue-specific (TS) enhancer drives the expression of Cas9. Coding exons of a sample gene of interest are in orange and untranslated regions are in grey. Two ubiquitously expressed gRNAs (gRNA1 and gRNA2) target different sites in the coding sequence. The Cas9 creates double strand breaks which can result in either indels (red bars) or a large deletion between the target sites (dash line).

## RESULTS

### Generation and evaluation of tissue-specific Cas9 lines

Our CRISPR-TRiM strategy (Fig. 1) relies on the availability of efficient tissue-specific Cas9 transgenes. To simplify the generation of tissue-specific Cas9 lines, we developed a Cas9 Gateway destination vector pDEST-APIC-Cas9 using the pAPIC (attB P-element insulated CaSpeR) backbone^23^ (Fig. 2a). Tissue-specific enhancers can be conveniently swapped into this vector through the Gateway LR reaction to generate Cas9-expression constructs. This cloning strategy is compatible with over 14,000 FlyLight^24^ and VT^25^ enhancers whose expression profiles for multiple developmental stages and tissues in *Drosophila* are publicly available. Several features of pDEST-APIC-Cas9 (Fig. 2a) make it optimal for making enhancer-driven Cas9 transgenes: (1) The Gypsy insulators flanking the Cas9 transgene boost transgene expression and reduce positional effects^26,27^. (2) A synthetic promoter combining the Hsp70 core promoter and additional motifs (Inr, MTE, and DPE)^28^ is predicted to support a wide range of enhancers and to offer higher expression than the widely used promoter DSCP^28^. (3) The combination of an intron in the 5’UTR and *His2Av* polyA further improves enhancer-driven transgene expression^23^. (4) Inclusion of both attB and P-element sequences provides flexibility for making transgenes either through random insertions or by targeted integration^23,29^.

**Fig. 2.**
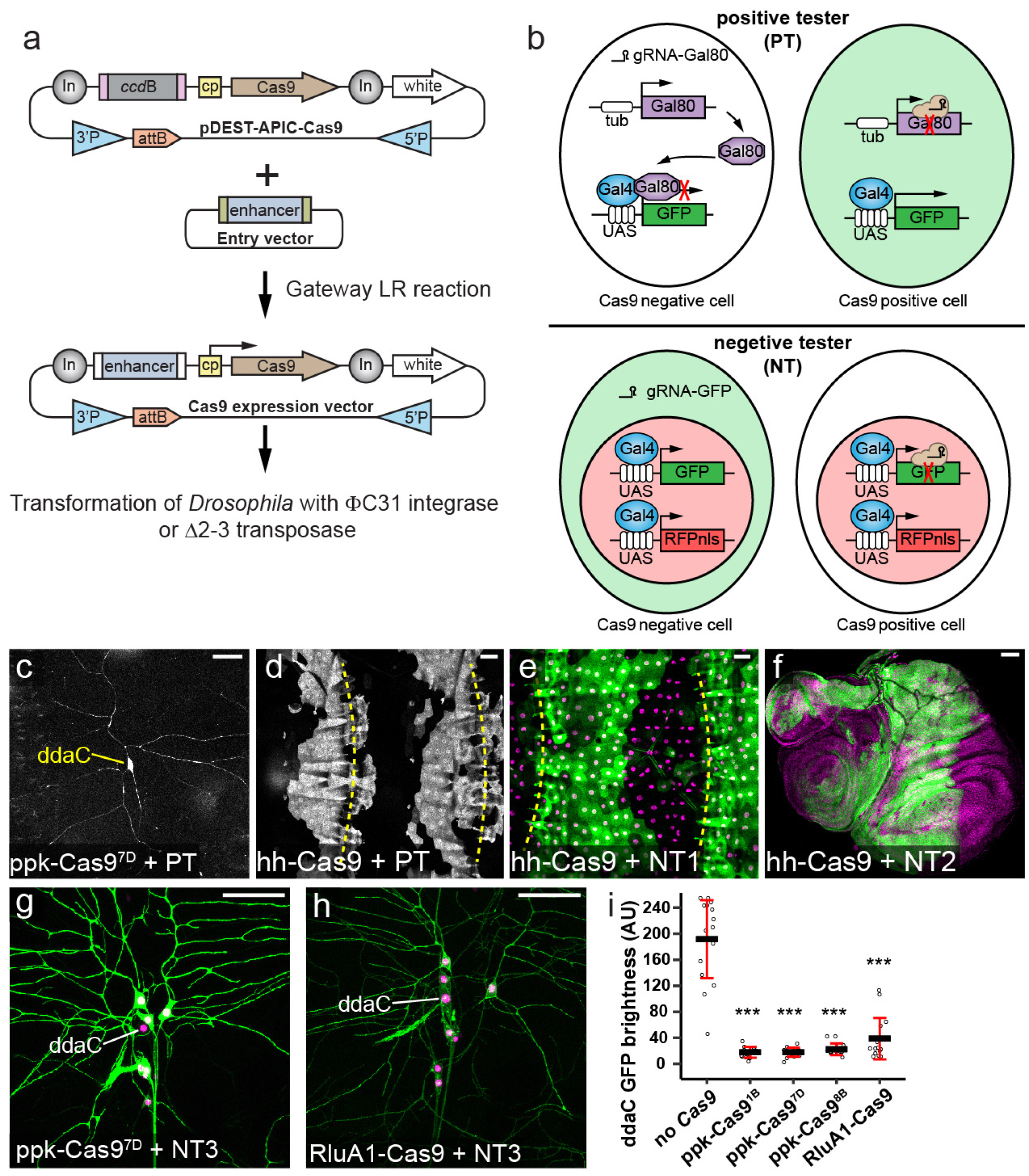
Generation and evaluation of tissue-specific Cas9 lines. (a) Diagram of Gateway cloning and transgenesis of Cas9 expression vectors. In, Gypsy insulator; cp, core promoter; 3’P and 5’P, P-element sequences. (b)Diagramsof positivetester (PT)and negative tester (NT), illustrating how Cas9-expressing cells are visualized by each type of tester. In PT, two ubiquitous gRNAs target Gal80. InNT, two ubiquitous gRNAs target GFP. Full genotypes of Cas9 testers are in Table S1.(c and d) Patterns of Cas9 activity in *ppk-Cas9*^*7D*^ (c) and *hh-Cas9* (d) as visualized by PT. (e and f) Patterns of *hh-Cas9* activity in the larval epidermis as visualized by NT1 (e) and inwing, haltere, and leg imaginal discs as visualized byNT2 (f). The positions of body wall segmental borders (muscle attachment sites) are indicated by yellow broken lines in (d) and (e). *ppk-Cas9* is predicted to be active in C4da neurons, including ddaC. *hh-Cas9* is predictedto be active in the posterior compartments of epidermal segments and imaginal discs. (g and h) Patterns of Cas9 activity in *ppk-Cas9*^*7D*^ (g) and *RluA1-Cas9* (h) as visualized by NT3. The cell bodies of ddaC neurons are indicated. (i) Quantification of ddaC GFP brightness in NT3 crosses usingcontrol (noCas9) and various da neuron-specific Cas9 lines. ***p≤0.001; one-way ANOVA and Dunnett’s test. (F=35.23, df=4). (p<2e-16 for *ppk-Cas9*^*1B*^; p<2e-16 for *ppk-Cas9*^*7D*^; p<2e-16 for *ppk-Cas9*^*8B*^; p<2e-16 for *RluA1-Cas9*). n=16 neurons for each genotype. Black bar, mean; red bars, SD. Scale bars, 50 μm)

An ideal tissue-specific Cas9 should be consistently and robustly expressed in the tissue of interest but not in unintended tissues. In practice, the insertion site in the genome often modifies the expression pattern, timing, and level of a transgene^30^. This position effect could impact the tissue-specificity and efficiency of mutagenesis. To evaluate Cas9 transgenes, we developed a series of tester lines, with the positive tester positively labeling Cas9-expressing cells and negative testers negatively labeling Cas9-expressing cells (Table S1). The positive tester ubiquitously expresses Gal80, Gal4, and two gRNAs targeting Gal80; and it also contains a UAS-driven GFP (Fig. 2b). In cells that do not express Cas9, Gal80 suppresses Gal4 activity, thereby inhibiting GFP expression. In contrast, in Cas9-expressing cells, the gRNAs induce mutations in Gal80 and thus allow Gal4-driven GFP expression. As examples, we generated random insertions of *ppk-Cas9* and *hh-Cas9* and evaluated their tissue specificities using the positive tester. The *ppk* enhancer is specific to class IV dendritic arborization (C4da) sensory neurons growing on the larval body wall^31^, while the R28E04 enhancer of *hh* drives epidermal expression in the posterior half of every hemisegment (http://flweb.janelia.org). The positive tester allowed us to identify the *ppk-Cas9* and *hh-Cas9* insertions that most resemble the expected patterns (Fig. 2c and d).

Negative testers help further evaluate the efficiency of Cas9 transgenes in inducing mutations. A negative tester contains a ubiquitous or tissue-specific Gal4, a UAS-driven cytosolic or membrane GFP, a UAS-driven nuclear RFP, and two ubiquitous gRNAs targeting GFP (Fig. 2b). With a negative tester, cells that do not express Cas9 are dually labeled by both GFP and the nuclear RFP. In contrast, Cas9-expressing cells are only labeled by the nuclear RFP, due to GFP mutagenesis. When crossed to negative testers ubiquitously expressing Gal4, *hh-Cas9* as expected caused loss of GFP in the posterior compartments of larval epidermal segments (Fig. 2e) and imaginal discs (Fig. 2f). A neuronal negative tester expressing the membrane marker CD8-GFP in all da neurons (NT3) showed that *ppk-Cas9* specifically knocked out GFP in C4da neurons (Fig. 2g). Negative testers are particularly useful for comparing the efficiency of Cas9 lines in mutagenesis: Lower persistent GFP signals likely reflect earlier-acting Cas9. Using NT3, we detected small but consistent differences among three efficient *ppk-Cas9* insertions (Fig. 2i), with two insertions (*ppk-Cas9*^*1B*^ and *ppk-Cas9*^*7D*^) outperforming the third one (*ppk-Cas9*^*8B*^). In comparison, Cas9 driven by a pan-da *RluA1* enhancer is less efficient in mutating GFP (Fig. 2h), leading to higher and variable levels of remaining GFP in C4da neurons (Fig. 2i).

The Cas9 Gateway cloning vector and the Cas9 tester lines together provide a convenient toolbox for generating and identifying Cas9 transgenes that are most efficient for CRISPR-TRiM.

### Optimization of multi-gRNA design for tissue-specific gene knockout in *Drosophila*

Being able to express multiple gRNAs from a single transgenic construct is desirable for CRISPR-TRiM, as more gRNAs can increase the chance of LOF in a single gene and also enable simultaneous mutagenesis of multiple genes^17–19^. A common strategy for making multiplex gRNA constructs in *Drosophila* is to use two or three ubiquitous U6 promotors in tandem, each driving a gRNA separately^18^. For this purpose, U6:1 and U6:3 promoters have been found to drive high gRNA expression^18^.

Alternatively, polycistronic gRNA designs with intervening tRNA sequences have also been reported to be effective in expressing multiple gRNAs in plants and *Drosophila*^19,32^. We wished to optimize the multi-gRNA strategy to achieve the greatest mutagenic efficiency in somatic cells. We thus compared four dual-gRNA designs that carry the same two targeting sequences for GFP (Fig. 3a). These constructs were made with a P-element/attB dual transformation vector that uses *mini-white* as the selection marker. Three of them (forward, reverse, and insulated) are variants of a U6:1-gRNA-U6:3-gRNA strategy described previously^18^, with differences in the orientation of the gRNA cassette and the use of an insulator to separate *mini-white* and the gRNAs. Our fourth design (tgFE) builds upon the tRNA-gRNA strategy^19^ and introduces an A-U base pair flip and an extension of the Cas9-binding hairpin (F+E modifications) in the gRNA scaffold^22^, which have been shown to greatly improve the targeting of Cas9 to the genomic DNA.

**Fig. 3.**
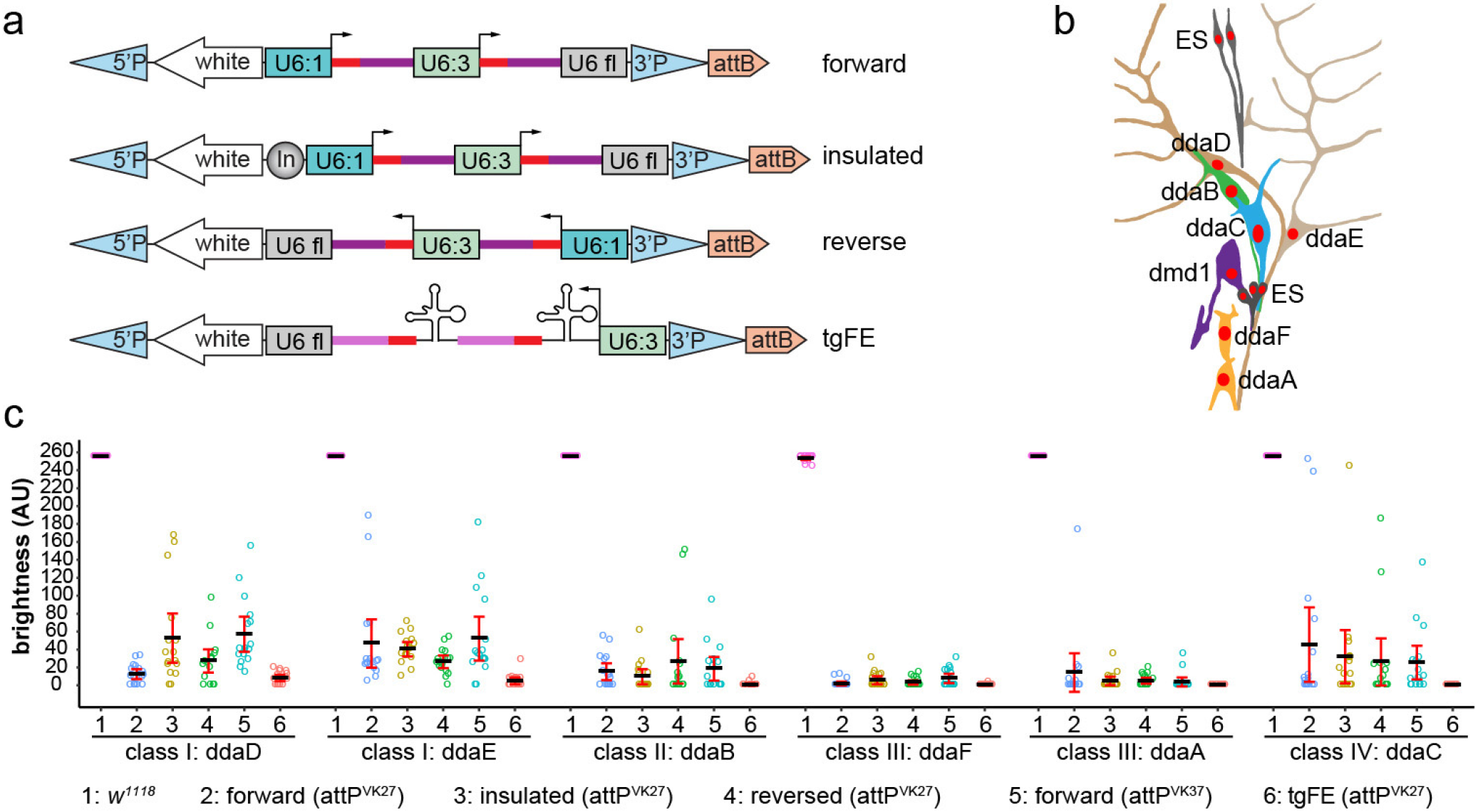
Optimization of multi-gRNA design for tissue-specific gene knockoutin *Drosophila*. (a) Four designs of multi-gRNA transgenic vectors. U6:1 and U6:3, U6 promoters; U6fl, U6 3’ flanking sequence; In, Gypsy insulator. Red bars, gRNA targeting sequence; dark magenta bars, original gRNA scaffold; light magenta bars, E+F gRNA scaffold. (b) Diagram of the dorsal cluster of larval peripheral sensory neurons. (c) Comparison of a control (1) and various *gRNA-GFP* lines in eliminating GFP signal in each dorsal da neuron using *RluA1-Cas9*. Daneurons express *UAS-CD8-GFP* driven by *nsyb-Gal4*. The integration site for each gRNA line is indicated in the parenthesis. The GFP signals in most control neurons are saturated under the setting used. Each circle represents an individual neuron (n=16 for each column). Black bar, mean; red bars, SD.

To compare these constructs, we used the larval PNS to evaluate the efficiency of GFP knockout in individual neurons. The dorsal cluster of sensory neurons in every abdominal segment contains 6 da neurons belonging to 4 classes^33^ (Fig. 3b), allowing for accurate measurement of fluorescence intensity at single cell resolutions. To detect sensitively differences in gRNA efficiency, we used the weakly expressed and ineffective *RluA1-Cas9* (Fig. 2h, i) to knock out *UAS-CD8-GFP* driven by pan-neural *nsyb-Gal4*^34^. We found that the performance of gRNAs based on U6:1-gRNA-U6:3-gRNA varied greatly for the same neuron and that none of these designs are efficient enough to remove GFP in all neurons (Fig. 3c). In contrast, the tgFE design was far superior, with near complete elimination of GFP signals in almost all neurons examined (Fig. 3c). Because previous studies indicated that the use of tRNA in polycistronic gRNAs does not seem by itself to enhance mutagenesis^19^, we think the tgFE design’s high efficiency is likely due to the F+E gRNA scaffold. An additional benefit of the tgFE strategy is the convenient cloning of 2-6 gRNAs in a single step.

### Efficiency of dual gRNA-mediated DNA deletion at the single cell level

When using two gRNAs to target the same gene, Cas9-mediated DSBs can result in indels at both target sites, or large DNA deletions between the two target sites^9^. Deletion of a larger piece of DNA is more likely to generate a null allele. To investigate the frequency of large deletions caused by two gRNAs in individual cells, we constructed a reporter *nSyb-tdGFP* (“td” standing for tandem dimer) (Fig. 4a) that labels all 12 neurons in the dorsal cluster of PNS sensory neurons (Fig. 3b, 4b). In addition, we designed 7 gRNAs (0 to 6) targeting different sites in the non-coding sequence of this reporter, with site 0 located before the nSyb enhancer, site 1 immediately after the enhancer, and the remaining sites at various distances downstream of the tdGFP coding sequence (Fig. 4a). We reasoned that small indels at any of these target sites would be unlikely to abolish GFP expression, but large deletions between site 0 and any of the other targeting sites would (Fig. 4c). As a control, we included a gRNA pair that targets two sites in the tdGFP coding sequence (gRNA-GFP) and therefore is predicted to remove GFP expression by either indels or large deletions.

**Fig. 4.**
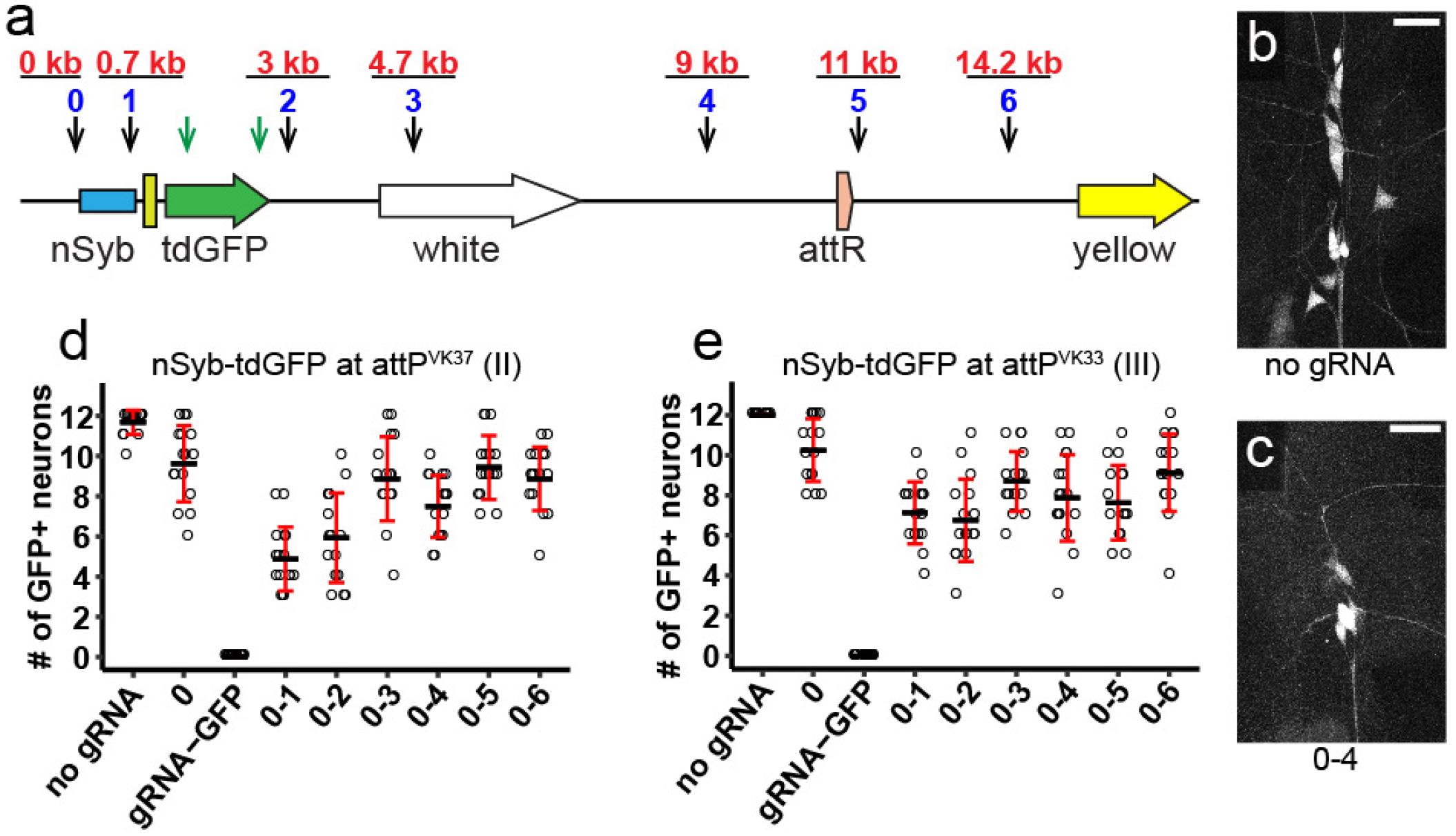
Efficiency of dual gRNA-mediated DNA deletion at the single cell level. (a) Diagram showing the *nSyb-tdGFP* reporterintegrated in the genome and gRNA target sites. Each blue number and the black arrow below it indicate a gRNA targeting non-coding sequence of the reporter. The distance of each gRNA from gRNA0 is indicated in red above the gRNA. The two green arrows indicate two gRNAs targeting the coding sequence of tdGFP. (b and c) Dorsal clusters of PNS neurons labeled by the reporter in a control animal (b) and an animal expressing gRNAs 0 and 4 (c). (d and e) Quantification of the number of GFP-positive neurons for each gRNA pair using *nSyb-tdGFP* inserted at *attP*^*VK37*^ (d) and *attP*^*VK33*^ (e) sites. Each circle represents an individual neuron (n=16 neurons for each genotype). Black bar, mean; red bars, SD.(Scale bars, 25 μm)

Using a strong and ubiquitously expressed *Act-Cas9*^18^, we tested the efficiencies of these gRNA pairs in eliminating GFP expression in individual neurons with two different *nSyb-tdGFP* insertions. In all animals examined, *gRNA-GFP* completely abolished GFP expression as expected (Fig. 4d, e), demonstrating the efficiency of DSB-mediated mutagenesis. Unexpectedly, gRNA 0 alone reduced numbers of labeled neurons in some animals (reduction mean±SD: 19.8%±15.7% for attP^VK37^ and 14.6%±13.1% for attP^VK33^) (Fig. 4d, e), likely due to deletions extending into regulatory elements in the nSyb enhancer. Pairing gRNA 0 with gRNAs 1-6 further reduced the number of labeled neurons (the range of reduction mean±SD for all pairs: 21.4%±13.3% to 59.4%±13.2% for attP^VK37^ and 24.0%±16.1% to 43.8%±17.1% for attP^VK33^), with a tendency for gRNA pairs positioned closer more often generating fewer GFP positive neurons (Fig. 4c-e). These data suggest that large deletions occur in random somatic cells and that an inverse correlation exists between deletion frequency and gRNA distance. Importantly, our results suggest that large deletions do not occur frequently enough to remove gene function in every cell such that indels in the coding region are more reliable for causing LOF.

### Enhancer-driven Cas9 is advantageous over Gal4-driven Cas9 for studying neuraldevelopment

Conditional mutagenesis can be achieved in *Drosophila* somatic cells using Gal4-driven Cas9^17–19^, but this method requires an intermediate transcription step that could potentially delay Cas9 expression.

Consistent with this assumption, *ppk-CD4-tdGFP* is expressed at least 8 hours earlier than *UAS-CD8-GFP* driven by *ppk-Gal4* in the embryo^23^. Thus, we predict that enhancer-driven Cas9 will result in earlier Cas9 action, thereby reducing perdurance of wildtype mRNA or protein products of the target gene made prior to mutation induction. We tested this hypothesis by comparing the effectiveness of enhancer-driven Cas9 and Gal4-driven Cas9 in knocking out CD4-tdGFP expression in C4da neurons (Fig. 5a). We observed more consistent and stronger reduction of GFP with *ppk-Cas9* insertions compared to *ppk-Gal4 UAS-Cas9* (*ppk>Cas9*) (Fig. 5b), although these differences were not statistically significant with our sample sizes. To ask whether even earlier Cas9 expression could lead to further GFP reduction, we made an enhancer-driven Cas9 that is expressed in sensory organ precursors (SOPs), the progenitor cells of da neurons^35^. Indeed, *SOP-Cas9* resulted in complete loss of GFP fluorescence in most animals (Fig. 5b).

**Fig. 5.**
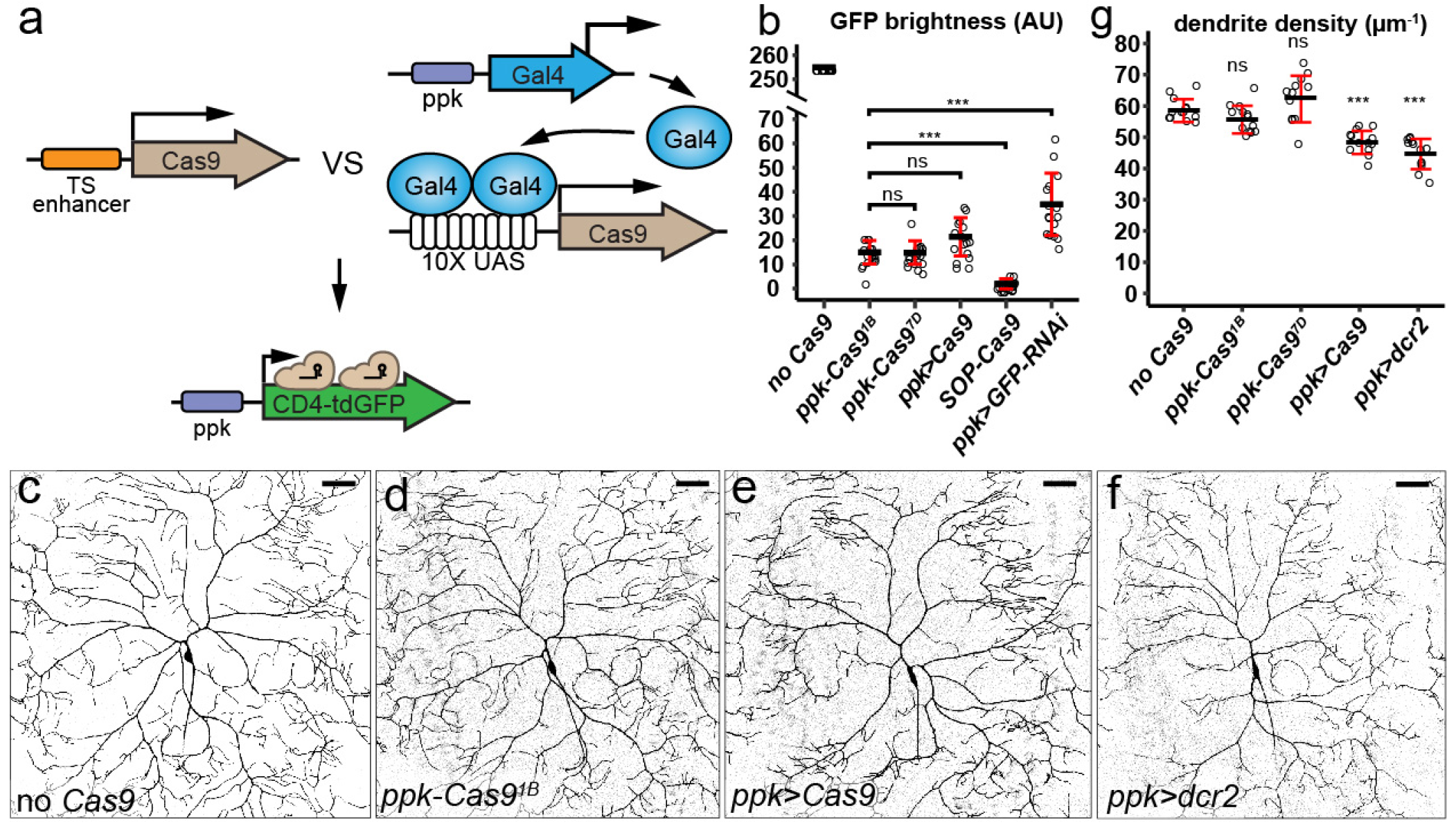
Enhancer-driven Cas9 is advantageous over Gal4-driven Cas9 instudyingneural development. (a) Diagram showing the comparison of tissue-specific (TS) enhancer-driven Cas9 and Gal4-driven Cas9 in knocking out *ppk-CD4-tdGFP*. (b) Quantification of GFP brightness in C4da neurons in the control, Cas9-expressing animals, and GFP knockdown animals. GFP signals in the control (no Cas9) are saturated. ***p≤0.001; ns, not significant; one-way ANOVA and Tukey’s HSD test (F=72.119, df=4). Exact p-values for comparison between *ppk-Cas9*^*1B*^ and *ppk>GFP-RNAi*, p<1e-04; *ppk-Cas9*^*1B*^ and *SOP-Cas9*, p<1e-04; *ppk-Cas9*^*1B*^ and *ppk>Cas9*, p=0.121; *ppk-Cas9*^*1B*^ and*ppk-Cas9*^*7D*^, p=1. (c-f) DdaC neurons in the control (c) and animals expressing *ppk-Cas9*^*1B*^ (d), *ppk-Gal4-*driven Cas9 (e), and ppk-*Gal4-*driven *dcr2* (f). (g) Quantification of dendrite density in genotypes indicated. ***p≤0.001; ns, not significant; one-way ANOVA and Dunnett’s test. (F=26.44,df=4) (p=0.386 for *ppk-Cas9*^*1B*^; p=0.197 for *ppk-Cas9*^*7D*^; p<0.001 for *ppk>Cas9*; p<0.001 for *ppk>dcr2*). Each circle represents an individual neuron. n=16 for genotypes in (b) and n=13 for genotypes in (g). Black bar, mean; red bars, SD. (Scale bars, 50 μm)

High levels of Cas9 have been reported to be cytotoxic^20^. Consistent with our supposition that Gal4-driven Cas9 generally produces more Cas9 protein than enhancer-fusion versions, we found that *ppk>Cas9* caused obvious dendrite reduction in C4da neurons even in the absence of gRNAs while *ppk-Cas9* lines had much weaker impacts on dendrite morphology (Fig. 5c-e, g). These data suggest that high levels of Cas9 in post-mitotic neurons are not desirable for studying neuronal morphogenesis and that enhancer-driven Cas9 could alleviate this concern.

We also compared the effects of RNAi-mediated suppression of GFP expression and CRISPR/Cas9-induced GFP mutagenesis. CD4-tdGFP was knocked down with a publicly available *UAS-GFP-RNAi* line^36^ driven by *ppk-Gal4*. We also co-expressed Dicer-2 (Dcr-2) in C4da neurons to enhance double strand RNA (dsRNA)-mediated mRNA knockdown^2^. RNAi was found to be less efficient in eliminating GFP than CRISPR-mediated mutagenesis by either enhancer-driven Cas9 or Gal4-drivenCas9 (Fig. 5b). In addition, we found that Dcr-2 overexpression, which is commonly employed in *Drosophila* RNAi experiments, caused an even stronger dendrite reduction than *ppk>Cas9* (Fig. 3f, g), indicating that Dcr-2 also has cytotoxicity in neurons.

Our results suggest that enhancer-driven Cas9 outperforms Gal4-driven Cas9 in tissue-specific mutagenesis and that the CRISPR-TRiM method is more effective than RNAi in LOF studies.

### Post-mitotic knockout of *Ptp69D* reveals its function in C4da neurons

To validate the effectiveness of CRISPR-TRiM in studying neuronal morphogenesis, we knocked out the receptor protein tyrosine phosphatase *Ptp69D* in C4da neurons. Using hemizygous *Ptp69D* mutants and MARCM, we previously found that loss of *Ptp69D* in C4da neurons caused dendritic reduction with shortened terminal dendrites^37^. As hemizygous mutants completely lack zygotic transcription and MARCM removes the *Ptp69D* gene before the birth of neurons, it is unclear from our previous results whether mutagenesis after the birth of neurons (post-mitotic mutagenesis) would be sufficient to remove Ptp69D function. We thus made a *gRNA-Ptp69D* line expressing three gRNAs, each targeting a distinct site in the *Ptp69D* coding sequence.

To validate the efficiency of *gRNA-Ptp69D*, we established a “Cas9-LEThAL” (for <u>Cas9-</u>induced lethal <u>e</u>ffect <u>th</u>rough the <u>a</u>bsence of <u>L</u>ig4) assay (Fig. S2) that was adapted from a previously described method for assessing injection-based gRNA efficiency^38^. Efficient gRNAs for non-essential genes, such as a published gRNA for *e*^18^ (Table S2), cause male-specific lethality in pupal stages when males carrying gRNAs are crossed to *Act-Cas9 lig4* homozygous females. But if the target gene is essential, in the same cross, efficient gRNAs should cause lethality of both males and females similar to homozygous mutants. *gRNA-Ptp69D* caused all animals to die at late pupal stages in this assay, indicating that this gRNA line is efficient (Table S2).

We knocked out *Ptp69D* in C4da neurons using both *ppk-Cas9* and *SOP-Cas9*. As the *ppk* enhancer only becomes active in stage-16 embryos after the birth of C4da neurons^31^, *ppk-Cas9* would only induce mutations post-mitotically. In contrast, the *SOP* enhancer turns on in sensory organ precursors^39^ that divide twice to give rise to da neurons^40^, enabling *SOP-Cas9* to act before the neuronal birth. We found that both *ppk-Cas9* and *SOP-Cas9* caused consistent and similar degrees of dendritic reduction in C4da neurons in late 3^rd^ instar larvae (Fig. 6a-d). In both cases, the extent of dendrite reduction caused by CRISPR-TRiM were also similar to that in *Ptp69D^14^/ Df(3L)8ex34* hemizygous null mutant larvae^37^ (Fig. 6e, f). These data suggest that post-mitotic mutagenesis is sufficient to remove *Ptp69D* gene function, which is consistent with Ptp69D being a neuronal type-specific transmembrane receptor^37,41^. Our results demonstrate that CRISPR-TRiM can efficiently eliminate gene function in a tissue-specific manner.

**Fig. 6.**
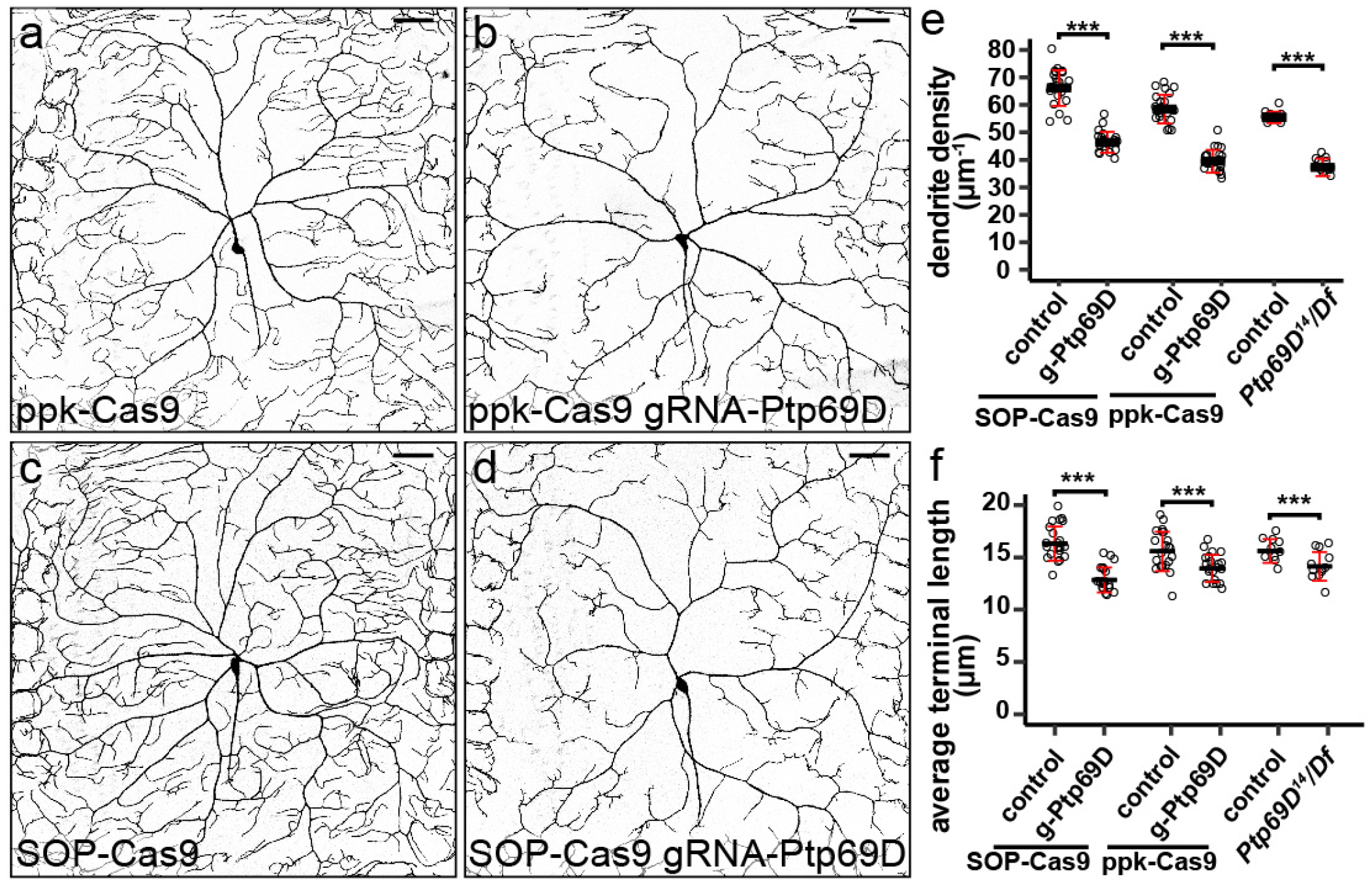
Post-mitotic knockout of *Ptp69D* is sufficient to reveal its function in C4da neurons. (a-d) DdaC neurons in *ppk-Cas9* control (a), *ppk-Cas9 gRNA-Ptp69D* (b), *SOP-Cas9* control (c), and *SOP-Cas9 gRNA-Ptp69D* (d). (e and f) Quantification of total dendrite density (e) and average terminal dendrite length (f) in the genotypes indicated. Each circle represents an individual neuron: *ppk-Cas9* (n=22); *ppk-Cas9 gRNA-Ptp69D* (n=21); *SOP-Cas9* (n=22); *SOP-Cas9 gRNA-Ptp69D* (n=20). Data for *Ptp69D^14^/Df(3L)8ex34* (n=12) and its control (n=10) are cited from^37^ for comparison. ***p≤0.001; Unpaired t-test. For e, *SOP-Cas9 gRNA-Ptp69D*, p<0.0001, t=11.84, df=40; *ppk-Cas9 gRNA-Ptp69D*, p<0.0001, t=13.14, df=41; *Ptp69D^14^/Df(3L)8ex34*, p<0.0001, t=10.39, df=20. For f, *SOP-Cas9 gRNA-Ptp69D*, p<0.0001, t=7.79, df=40; *ppk-Cas9 gRNA-Ptp69D*, p=0.0013, t=3.46, df=41; *Ptp69D^14^/Df(3L)8ex34*, p=0.128, t=2.73, df=20. Black bar, mean; redbars, SD. (Scale bars, 50 μm)

### CRISPR-TRiM reveals the redundancy and perdurance of NSF and SNAP.b genes in dendrite morphogenesis

In *Drosophila*, CRISPR/Cas9 has been used to simultaneously mutagenize multiple genes in somatic cells19. Such an application would be very useful for studying the roles of redundant genes during development. To test whether CRISPR-TRiM can efficiently knock out multiple genes that may exhibit redundant functions, we targeted SNARE complex components in C4da neurons. Because SNAREs are required for all vesicle fusions^42^, interference with the complex should severely hamper C4da dendrite growth. *Drosophila* contains two NSF genes (*comt/Nsf1* and *Nsf2*) which are necessary for the recycling of the SNARE complex after membrane fusion^43^. *Drosophila* also has three SNAP-25 paralogues (*Snap24*, *Snap25*, and *Snap29*) that encode the SNAP.b (or Qb.IV) group of SNARE proteins thought to be involved in secretion^44^. The potential functional redundancy of NSF and SNAP.b genes has not to date been examined during neuronal morphogenesis.

To conduct CRISPR-TRiM analyses, we used the tgFE design to generate dual-gRNA constructs for every NSF and SNAP.b gene. Also using the tgFE design, we made 4-gRNA constructs to knock out *Nsf1*/*Nsf2* simultaneously and *Snap24*/*Snap25* simultaneously, and a 6-gRNA construct to knock out all three SNAP.b genes. The efficiencies of these gRNA lines were first validated with the Cas9-LEThAL assay (Table S2). The lethal phase induced by each single-gene gRNA line was consistent with published results for null mutants of the corresponding gene, indicating that the gRNAs are efficient in mutagenesis. We found that *gRNA-Nsf1-Nsf2* was as effective as *gRNA-Nsf2* in causing lethality at 1^st^ instar larvae, while *gRNA-Nsf1* caused lethality in late pupae. This suggests that increasing the number of gRNAs from 2 to 4 in one construct does not reduce the efficiency of gRNAs. Interestingly, compared to *gRNA-Snap24* or *gRNA-Snap25* alone, which produced animals surviving to the late pupal stage, *gRNA-Snap24-Snap25* caused lethality at the 1st instar larvae, demonstrating that Snap24 and Snap25 are redundantly required for the larval development. *gRNA-Snap24-Snap25-Snap29* further advanced the lethal phase to late embryos, suggesting that Snap29 is also redundant with Snap24 and Snap25.

We knocked out NSF and SNAP.b genes in C4da neurons using both *ppk-Cas9* and *SOP-Cas9*. Removing individual NSF genes did not cause obvious dendritic reductions (Fig. S1a-c, h-j), but surprisingly *SOP-Cas9/gRNA-Nsf2* neurons instead showed a mild but consistent increase of dendrite length and density (Fig. 7g-h,). Knocking out both *Nsf1* and *Nsf2* using *ppk-Cas9* produced weak and variable C4da dendrite reduction (Fig. 7a, b, g-h), while *SOP-Cas9/gRNA-Nsf1-Nsf2* animals showed stronger and more consistent dendrite reductions (Fig. 7e, g-h). These data suggest that *Nsf1* and *Nsf2* act redundantly to promote dendrite growth. Furthermore, the observation that *SOP-Cas9* caused a stronger phenotype than *ppk-Cas9* suggest that NSF gene products made in sensory organ precursors are not sufficient to support dendrite growth but those made at the time of post-mitotic mutagenesis allow neurons to grow a significant amount of dendrites.

**Fig. 7.**
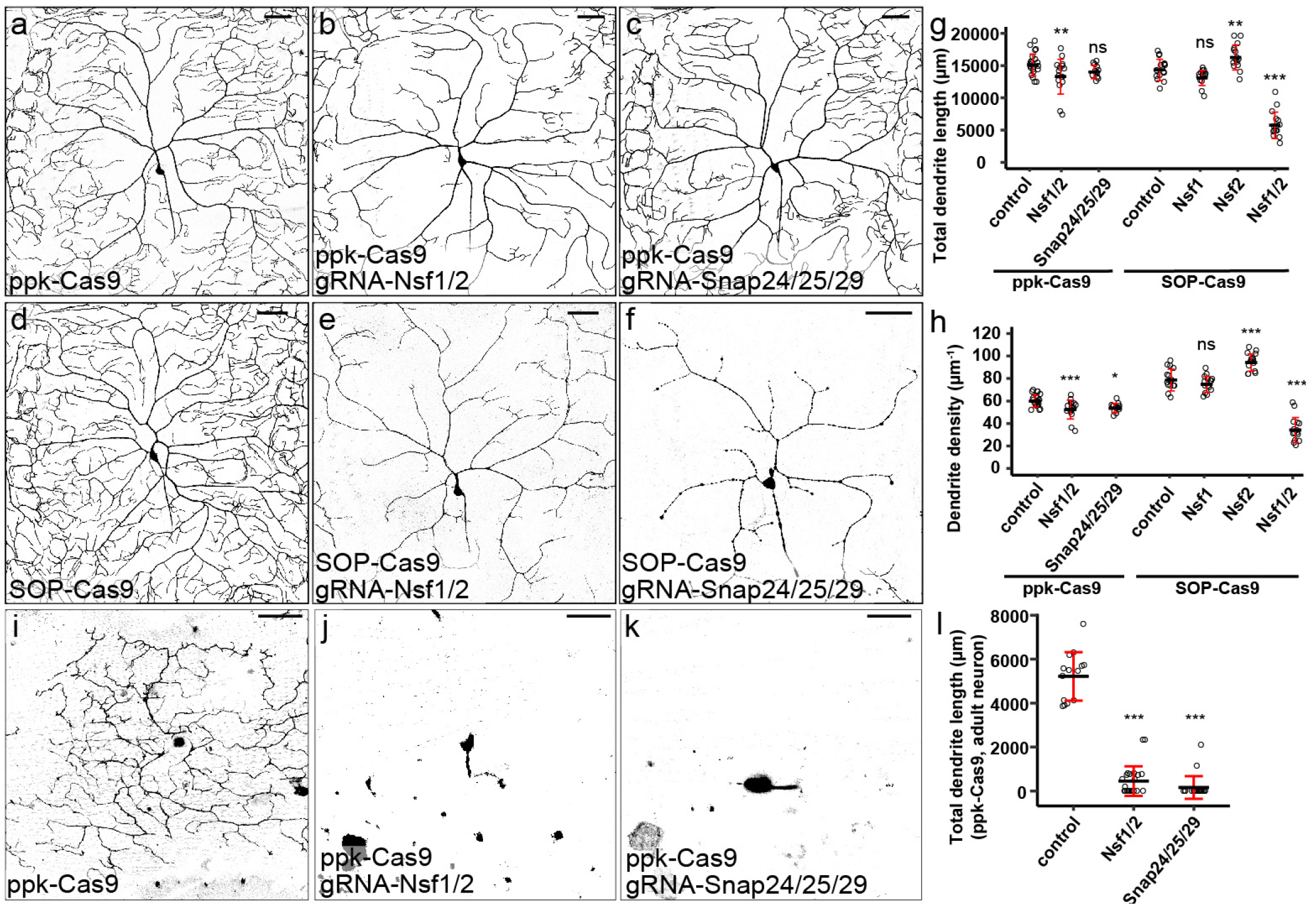
CRISPR-TRiM analyses of NSF and SNAP.b genes in C4da dendrite morphogenesis. (a-c) DdaC neurons in *ppk-Cas9* (a), *ppk-Cas9 gRNA-Nsf1-Nsf2* (b), and *ppk-Cas9 gRNA-Snap24-Snap25-Snap29* (c). (d-f) DdaC neurons in *SOP-Cas9* (d), *SOP-Cas9 gRNA-Nsf1-Nsf2* (e), and *SOP-Cas9 gRNA-Snap24-Snap25-Snap29* (f). (g and h) Quantification of total dendrite length (g) and dendrite density (h) in the genotypes indicated. Each circle represents an individual neuron: *ppk-Cas9* (n=22); *ppk-Cas9 gRNA-Nsf1-Nsf2* (n=16); *ppk-Cas9 gRNA-Snap24-Snap25-Snap29* (n=11); *SOP-Cas9* (n=15), *SOP-Cas9 gRNA-Nsf1* (n=15); *SOP-Cas9 gRNA-Nsf2* (n=15); *SOP-Cas9 gRNA-Nsf1-Nsf2* (n=15). *p≤0.05; **p≤0.01; ***p≤0.001; ns, not significant; one-way ANOVA and Dunnett’s test. For g, F=103.49, df=6; p=0.0151 for *ppk-Cas9 gRNA-Nsf1-Nsf2*; p=0.263 for *ppk-Cas9 Snap24-Snap25-Snap29*; p=0.138 for *SOP-Cas9 gRNA-Nsf1*; p=0.01 for *SOP-Cas9 gRNA-Nsf2*; p<0.001 for*SOP-Cas9 gRNA-Nsf1-gRNA2*. For h, F=44.016, df=6; p=0.00161 for *ppk-Cas9 gRNA-Nsf1-Nsf2*; p=0.0211 for *ppk-Cas9 Snap24-Snap25-Snap29*; p=0.506 for *SOP-Cas9 gRNA-Nsf1*; p<1e-4 for *SOP-Cas9 gRNA-Nsf2*; p<1e-4 for *SOP-Cas9 gRNA-Nsf1-gRNA2*. (i-k) V’ada neurons in day 0 adults of *ppk-Cas9* (i), *ppk-Cas9 gRNA-Nsf1-Nsf2* (j), *ppk-Cas9 gRNA-Snap24-Snap25-Snap29* (k). (l) Quantificationof total dendrite length of adult v’ada neurons expressing *ppk-Cas9* and the gRNAs indicated. Each circle represents an individual neuron: *ppk-Cas9* (n=14), *ppk-Cas9 gRNA-Nsf1-Nsf2* (n=23), *ppk-Cas9 gRNA-Snap24-Snap25-Snap29* (n=20). ***p≤0.001; one-way ANOVA and Dunnett’s test (F=220.87,df=2). p<1e-10 for *gRNA-Nsf1-Nsf2*; p<1e-10 for *gRNA-Snap24-Snap25-Snap29*. Black bar, mean; red bars, SD. (Scale bars, 50 μm)

Tissue-specific knockout of individual SNAP.b genes using *ppk-Cas9* or *SOP-Cas9* produced either noobvious phenotypes (for *Snap24* and *Snap25*) or weak dendrite reductions (for *Snap29*) (Fig. S1d-f, k-m). Knocking out both *Snap24* and *Snap25* similarly did not cause obvious dendrite defects (Fig. S1g, n), which is somewhat surprising considering that *Snap24* and *Snap25* play redundant roles in synaptic transmission at neuromuscular junctions (NMJs)^45^. We next knocked out all three SNAP.b genes using *gRNA-Snap24-Snap25-Snap29*. While *ppk-Cas9-*mediated knockout only slightly reduced dendrite density (Fig. 7c, g-h), *SOP-Cas9*-mediated knockout caused strong C4da dendrite reduction and degeneration (n=16/19neurons) in second instar larvae (Fig. 7f) and late larval lethality. Although *SOP-Cas9* is highly efficient in da neurons as shown by the NT3 negative tester (Fig. S1o), this lethality might be independent of neuronal defects, because *SOP-Cas9* also labeled a small number of random larval epidermal cells with the positive tester (Fig. S1p). Nevertheless, our results suggest that, like NSF genes, all three SNAP.b genes are redundantly required in C4da neurons and that mutagenesis before the neuronal birth is required to unmask the LOF phenotype of SNAP.b genes.

As the SNARE machinery is required for all vesicle trafficking in the cell, we were curious why
knocking out all NSF or all SNAP.b genes in neurons with *SOP-Cas9* was not sufficient to suppress all dendritic growth. One possibility is that membrane trafficking-independent mechanisms exist that allow neurons to elaborate dendrites. Alternatively, NSF and SNAP.b gene products contributed maternally or made before *SOP-Cas9* activity persist long enough to support a small degree of larval dendrite growth. To distinguish between these possibilities, we turned to adult C4da neurons. C4da neurons ddaC and v’ada prune all their dendrites during metamorphosis and regrow new dendritic arbors in late pupae^46^.

Because dendritic pruning removes all existing gene products except for the residual amounts left in the cell body, dendrite regrowth must rely on new transcription. If NSF and SNAP.b genes are required for all dendrite growth, knocking out all NSF or SNAP.b genes during larval stages should prevent dendrite regrowth. Indeed, adult v’ada neurons lacking *Nsf1 Nsf2* or *Snap24 Snap25 Snap29* via *ppk-Cas9*-mediated knockout either did not regrow primary branches or showed severe reduction in total dendrite length (Fig. 7i-l). These data suggest that *ppk-Cas9* can effectively remove redundant genes in post-mitotic neurons and that neuronal dendrite growth absolutely requires SNARE function.

## DISCUSSION

In this study, we describe an optimized strategy we call CRISPR-TRiM for tissue-specific gene mutagenesis using CRISPR/Cas9 in *Drosophila*. To implement this method, we developed a toolkit for generating and evaluating enhancer-driven Cas9 lines, created convenient cloning vectors for making efficient multi-gRNA transgenes, and established an assay for assessing the mutagenic efficiency of transgenic gRNAs. Using our CRISPR-TRiM tools, we demonstrate that post-mitotic knockout of *Ptp69D* causes efficient LOF in neurons, while SNARE complex components are strongly redundant and perdurant in supporting neuronal dendrite development.

### Comparison of CRISPR-TRiM with other tissue-specific LOF methods

Flp/FRT-based mosaic analyses have been widely used for investigating tissue-specific roles of genes in *Drosophila*^47^. Among these techniques, MARCM and its variants are considered gold standards for neuronal studies due to the positive labeling of homozygous mutant cells and the single cell resolution they offer^1,48^. However, MARCM and other Flp/FRT-based mosaic analyses also have some obvious limitations. First, they require preexisting mutations in the gene of interest recombined with FRT on the appropriate chromosome arm. Second, because these techniques rely on mitotic chromosome crossovers which would result in wildtype “twin spots”, it is impossible to remove gene function in every cell of the tissue of interest. Third, these techniques require at least five genetic components in the final genotype, making it harder to introduce additional components. Lastly, generating cells mutant for multiple genes located on different chromosome arms is extremely difficult, if not impossible. In contrast, the bipartite CRISPR-TRiM system requires only transgenic components that are independent of all existing binary expression systems. Using efficient Cas9 and gRNA reagents, LOF in all cells of the target tissue can be expected. These features make CRISPR-TRiM much more convenient than traditional mosaic-based methods.

Compared to *UAS-Cas9* driven by tissue-specific Gal4s, our CRISPR-TRiM system has the advantages of faster Cas9 expression (and therefore more complete LOF) and decreased cytotoxicity due to lower Cas9 expression levels. An additional benefit of using enhancer-driven Cas9 in CRISPR-TRiM is that the Gal4/UAS system is available for other genetic manipulations in the same experiment.

In the last decade, several genome-wide *UAS-RNAi* resources have greatly accelerated gene identification and characterization in *Drosophila*^2,3^. However, RNAi results in incomplete LOF and suffers from off-target effects^4^. In comparison, CRISPR methods can generate true gene knockout, and ever-improving gRNA selection algorisms have mostly mitigated the off-target effects^49–51^. In addition, *UAS-Dcr-2* overexpression, which is often necessary for maximizing the knockdown efficiency of dsRNAs, can also cause deleterious effects in the expressing cells. The CRISPR-TRiM method can avoid most of these concerns.

### Caveats of CRISPR-TRiM and potential solutions

Due to the nature of CRISPR/Cas9-induced mutagenesis, CRISPR-TRiM will generate tissues composed of heterogenous cells carrying different mutations. This mosaicism could complicate phenotypic analysis, given that different mutations could impact gene function in diverse ways. Although immunostaining could alleviate this problem by revealing whether individual cells make the final protein product, antibodies are not always available nor are all assays compatible with immunostaining. For this reason, we recommend the use of at least two gRNAs for each target gene, because two gRNAs dramatically enhance the chance of mutagenesis.

Nonetheless, even with multiple efficient gRNAs we observed that CRISPR-TRiM sometimes produced variable phenotypes that varied from cell to cell (e.g. *Nsf1 Nsf2* knockout by *ppk-Cas9,* Fig. 7g, l), likely due to the differences in the timing of mutagenesis and/or the nature of the mutations induced in different cells. This variability could actually be beneficial for the analysis of tissues like da neurons where each cell can be evaluated separately: The mosaicism could reveal a fuller spectrum of phenotypes associated with different strengths of LOF. Moderately efficient gRNAs could potentially be exploited for this purpose.

### Designing efficient gRNA constructs and assessing gRNA efficiency

Our comparison of several dual-gRNA designs using the same targeting sequences revealed that the tgFE design is particularly efficient for mutagenesis in larval sensory neurons. We found that the same design also performs well in other somatic tissues (data not shown). The tgFE design combines tRNA-processing for releasing multiple gRNAs from a single transcript^19,32^ and F+E modifications that consist of an A-U flip and a hairpin extension in the gRNA scaffold^22^. We consider this design the most significant improvement for our CRISPR-TRiM system. This design may also work well in the germ line for creating heritable mutations.

Although large deletions induced by two gRNAs would be more effective in causing LOF, we found that the frequency of large deletions in somatic cells is too low to be reliable. Therefore, to maximize the chance of LOF mutagenesis, we recommend selecting targeting sites in coding sequences shared by all protein isoforms, preferably in conserved protein domains. In our experience, choosing common top hits by using multiple experimentally-validated gRNA selection algorithms^49,52,53^ usually yield very efficient gRNAs.

We also recommend evaluating the *in vivo* efficiency of gRNA lines using the Cas9-LEThAL assay before conducting CRISPR-TRiM analyses. In our hands, the lethal phase of male progeny in this assay reliably indicates gRNA efficiency for our CRISPR-TRiM experiments.

### Using CRISPR-TRiM to study neuronal morphogenesis

We provide two examples of CRISPR-TRiM analysis in C4da neurons. Our results show that the timing of mutagenesis and the perdurance of gene products influence the extent of LOF; therefore, these parameters must be considered when choosing the most appropriate *Cas9* line. The CRISPR-TRiM analysis of *Ptp69D* shows that post-mitotic mutagenesis is sufficient to cause its LOF, because Ptp69D either is expressed late in neuronal development or turns over quickly. In contrast, SNARE components are made early in the neuronal lineage and are highly perdurant. The early-acting *SOP-Cas9* therefore is required to reveal SNARE LOF phenotypes in neurons. Moreover, we found that dendrite regrowth of adult C4da neurons provides an opportunity to unmask fully the requirements of SNARE components for dendrite morphogenesis. This technique should be useful for circumventing potential perdurance because gene products are removed by dendrite pruning prior to the regrowth. Our results imply that perdurance could be an underappreciated concern for studying development roles of housekeeping genes in any mutation-based LOF analysis.

Previous studies have shown that *Snap-24* and *Snap-25* act redundantly to promote NMJ neurotransmitter release^45^. Our data indicate that an additional redundant gene, *Snap-29*, is also involved in C4da dendrite branching morphogenesis. Our NSF LOF data demonstrate that Nsf1 and Nsf2 play redundant roles in C4da neurons, which is consistent with the previous finding that these two genes can substitute for each other in the nervous system^43^. Of interest, we observed a distinction between NSF and SNAP.b LOF phenotypes. Both with *SOP-Cas9* in the larva and *ppk-Cas9* in the adult, SNAP.b LOF appears to produce a more severe dendritic reduction than NSF LOF. We believe this distinction
reflects the different roles of these proteins in the SNARE machinery. Because NSFs are responsible for recycling the SNARE complex after membrane fusion, newly synthesized SNARE components can still mediate vesicle fusion in the absence of NSFs. In contrast, SNAP.b LOF causes secretion to stop completely, thereby generating a stronger phenotype.

In summary, we present an optimized system for generating tissue-specific gene knockout using CRISPR/Cas9. The tools we developed can be applied to address a broad range of developmental, cell biological, and physiological questions in *Drosophila*.

## METHODS

Methods are in the Supplementary Information.

## ACKNOWLEDGEMENTS

We thank Ying Peng, Yi Guo, and Bloomington Stock Center for fly stocks; Norbert Perrimon and Addgene for plasmids; Michael Goldberg, Mariana Wolfner, David Deitcher, Dion Dickman, and Quan Yuan for critical reading and suggestions on the manuscript. This work was supported by a Cornell Fellowship awarded to H.J.; a Cornell start-up fund and NIH grants (R01NS099125 and R21OD023824) awarded to C.H.

## AUTHOR CONTRIBUTIONS

CH and ARP designed the experiments. BW and YH conducted molecular cloning. ARP performed imaging and quantification. CH, ARP, MLS,and HJ built genetic reagents used in this study. KL, TO, RF, MG, and YQ screened Cas9 transgenic lines. CH and ARP wrote the manuscript.

## REFERENCE

1 Lee, T. & Luo L. Mosaic analysis with a repressible cell marker for studies of gene function in neuronal morphogenesis.Neuron 22, 451–461 (1999).

2 Dietzl G. et al. A genome-wide transgenic RNAi library for conditional gene inactivation in Drosophila. Nature 448, 151–156, doi:10.1038/nature05954 (2007).

3 Ni J. Q. et al. A genome-scale shRNA resource for transgenic RNAi in Drosophila. Nat Methods8, 405–407, doi:10.1038/nmeth.1592 (2011).

4 Ma Y., Creanga A., Lum L. & Beachy P. A. Prevalence of off-target effects in Drosophila RNA interference screens. Nature 443, 359–363, doi:10.1038/nature05179 (2006).

5 Jinek M. et al. A programmable dual-RNA-guided DNA endonuclease in adaptive bacterial immunity. Science 337, 816–821, doi:10.1126/science.1225829 (2012).

6 Bassett A. R., Tibbit C., Ponting C. P. & Liu J. L. Highly efficient targeted mutagenesis of Drosophila with the CRISPR/Cas9 system. Cell Rep 4, 220–228, doi:10.1016/j.celrep.2013.06.020 (2013).

7 Gratz S. J. et al. Genome engineering of Drosophila with the CRISPR RNA-guided Cas9 nuclease. Genetics 194, 1029–1035, doi:10.1534/genetics.113.152710 (2013).

8 Yu Z. et al. Highly efficient genome modifications mediated by CRISPR/Cas9 in Drosophila.Genetics 195, 289–291, doi:10.1534/genetics.113.153825 (2013).

9 Ren X. et al. Optimized gene editing technology for Drosophila melanogaster using germ line-specific Cas9. Proc Natl Acad Sci U S A 110, 19012–19017, doi:10.1073/pnas.1318481110 (2013).

10 Kondo S. & Ueda R. Highly improved gene targeting by germline-specific Cas9 expression in Drosophila. Genetics 195, 715–721, doi:10.1534/genetics.113.156737 (2013).

11 Sebo Z. L., Lee H. B., Peng Y. & Guo Y. A simplified and efficient germline-specific CRISPR/Cas9 system for Drosophila genomic engineering. Fly (Austin) 8, 52–57, doi:10.4161/fly.26828 (2014).

12 Gaj T., Gersbach C. A. & Barbas C. F., 3rd. ZFN, TALEN, and CRISPR/Cas-based methods for genome engineering. Trends Biotechnol 31, 397–405, doi:10.1016/j.tibtech.2013.04.004 (2013).

13 Gratz S. J. et al. Highly specific and efficient CRISPR/Cas9-catalyzed homology-directed repair in Drosophila. Genetics 196, 961–971, doi:10.1534/genetics.113.160713 (2014).

14 Xue Z. et al. Efficient gene knock-out and knock-in with transgenic Cas9 in Drosophila. G3 (Bethesda) 4, 925–929, doi:10.1534/g3.114.010496 (2014).

15 Ghosh S., Tibbit C. & Liu J. L.Effective knockdown of Drosophila long non-coding RNAs by CRISPR interference. Nucleic Acids Res 44, e84, doi:10.1093/nar/gkw063 (2016).

16 Ewen-Campen B. et al. Optimized strategy for in vivo Cas9-activation in Drosophila. Proc Natl Acad Sci U S A 114, 9409–9414, doi:10.1073/pnas.1707635114 (2017).

17 Xue Z. et al. CRISPR/Cas9 mediates efficient conditional mutagenesis in Drosophila. G3 (Bethesda) 4, 2167–2173, doi:10.1534/g3.114.014159 (2014).

18 Port F., Chen H. M., Lee T. & Bullock S. L. Optimized CRISPR/Cas tools for efficient germline and somatic genome engineering in Drosophila. Proc Natl Acad Sci U S A 111, E2967–2976, doi:10.1073/pnas.1405500111 (2014).

19 Port F. & Bullock S. L. Augmenting CRISPR applications in Drosophila with tRNA-flanked sgRNAs. Nat Methods 13, 852–854, doi:10.1038/nmeth.3972 (2016).

20 Jiang W., Brueggeman A. J., Horken K. M., Plucinak T. M. & Weeks D. P. Successful transient expression of Cas9 and single guide RNA genes in Chlamydomonas reinhardtii. Eukaryot Cell 13, 1465–1469, doi:10.1128/EC.00213-14 (2014).

21 Ren X. et al. Enhanced specificity and efficiency of the CRISPR/Cas9 system with optimized sgRNA parameters in Drosophila. Cell Rep 9, 1151–1162, doi:10.1016/j.celrep.2014.09.044(2014).

22 Chen B. et al. Dynamic imaging of genomic loci in living human cells by an optimized CRISPR/Cas system. Cell 155, 1479–1491, doi:10.1016/j.cell.2013.12.001 (2013).

23 Han C., Jan L. Y. & Jan Y. N. Enhancer-driven membrane markers for analysis of nonautonomous mechanisms reveal neuron-glia interactions in Drosophila. Proc Natl Acad Sci U S A 108, 9673–9678, doi:10.1073/pnas.1106386108 1106386108 [pii] (2011).

24 Jenett A. et al. A GAL4-driver line resource for Drosophila neurobiology.Cell Rep 2, 991–1001, doi:10.1016/j.celrep.2012.09.011 (2012).

25 Kvon E. Z. et al. Genome-scale functional characterization of Drosophila developmental enhancers in vivo. Nature 512, 91–95, doi:10.1038/nature13395 (2014).

26 Roseman R. R., Pirrotta V. & Geyer P. K. The su(Hw) protein insulates expression of the Drosophila melanogaster white gene from chromosomal position-effects. EMBO J 12, 435–442 (1993).

27 Markstein M., Pitsouli C., Villalta C., Celniker S. E. & Perrimon N. Exploiting position effects and the gypsy retrovirus insulator to engineer precisely expressed transgenes. Nat Genet 40, 476–483, doi:10.1038/ng.101 (2008).

28 Pfeiffer B. D. et al. Tools for neuroanatomy and neurogenetics in Drosophila. Proc Natl Acad Sci U S A 105, 9715–9720, doi:10.1073/pnas.0803697105 (2008).

29 Venken K. J., He Y., Hoskins R. A. & Bellen H. J. P[acman]: a BAC transgenic platform for targeted insertion of large DNA fragments in D. melanogaster. Science 314, 1747–1751, doi:10.1126/science.1134426 (2006).

30 Weiler K. S. & Wakimoto B. T. Heterochromatin and gene expression in Drosophila. Annu Rev Genet 29, 577–605, doi:10.1146/annurev.ge.29.120195.003045 (1995).

31 Grueber W. B., Ye B., Moore A. W., Jan L. Y. & Jan Y. N. Dendrites of distinct classes of Drosophila sensory neurons show different capacities for homotypic repulsion. Curr Biol 13, 618–626 (2003).

32 Xie K., Minkenberg B. & Yang Y. Boosting CRISPR/Cas9 multiplex editing capability with the endogenous tRNA-processing system. Proc Natl Acad Sci U S A 112, 3570–3575, doi:10.1073/pnas.1420294112 (2015).

33 Grueber W. B., Jan L. Y. & Jan Y. N. Tiling of the Drosophila epidermis by multidendritic sensory neurons. Development 129, 2867–2878 (2002).

34 Pfeiffer B. D., Truman J. W. & Rubin G. M. Using translational enhancers to increase transgene expression in Drosophila. Proc Natl Acad Sci U S A 109, 6626–6631, doi:10.1073/pnas.1204520109 (2012).

35 Powell L. M., Zur Lage P. I., Prentice D. R., Senthinathan B. & Jarman A. P. The proneural proteins Atonal and Scute regulate neural target genes through different E-box binding sites. Mol Cell Biol 24, 9517–9526, doi:10.1128/MCB.24.21.9517-9526.2004 (2004).

36 Pastor-Pareja J. C. & Xu T. Shaping cells and organs in Drosophila by opposing roles of fat body-secreted Collagen IV and perlecan. Dev Cell 21, 245–256, doi:10.1016/j.devcel.2011.06.026 (2011).

37 Poe A. R. et al. Dendritic space-filling requires a neuronal type-specific extracellular permissive signal in Drosophila. Proc Natl Acad Sci U S A 114, E8062–E8071, doi:10.1073/pnas.1707467114 (2017).

38 Lee H. B., Sebo Z. L., Peng Y. & Guo Y. An optimized TALEN application for mutagenesis and screening in Drosophila melanogaster.Cell Logist 5, e1023423, doi:10.1080/21592799.2015.1023423 (2015).

39 Culi J. & Modolell J. Proneural gene self-stimulation in neural precursors: an essential mechanism for sense organ development that is regulated by Notch signaling. Genes Dev 12, 2036–2047 (1998).

40 Lai E. C. & Orgogozo V. A hidden program in Drosophila peripheral neurogenesis revealed: fundamental principles underlying sensory organ diversity. Dev Biol 269, 1–17, doi:10.1016/j.ydbio.2004.01.032 (2004).

41 Desai C. J., Popova, E. & Zinn K. A Drosophila receptor tyrosine phosphatase expressed in the embryonic CNS and larval optic lobes is a member of the set of proteins bearing the “HRP” carbohydrate epitope. J Neurosci 14, 7272–7283 (1994).

42 Wickner W. & Schekman R. Membrane fusion. Nat Struct Mol Biol 15, 658–664 (2008).

43 Golby J. A., Tolar L. A. & Pallanck L. Partitioning of N-ethylmaleimide-sensitive fusion (NSF) protein function in Drosophila melanogaster: dNSF1 is required in the nervous system, and dNSF2 is required in mesoderm. Genetics 158, 265–278 (2001).

44 Kloepper T. H., Kienle C. N. & Fasshauer D. An elaborate classification of SNARE proteins sheds light on the conservation of the eukaryotic endomembrane system. Mol Biol Cell 18, 3463–3471, doi:10.1091/mbc.E07-03-0193 (2007).

45 Vilinsky I., Stewart B. A., Drummond J., Robinson I. & Deitcher D. L. A Drosophila SNAP-25 null mutant reveals context-dependent redundancy with SNAP-24 in neurotransmission. Genetics 162, 259–271 (2002).

46 Shimono K. et al. Multidendritic sensory neurons in the adult Drosophila abdomen: origins, dendritic morphology, and segment- and age-dependent programmed cell death. Neural Dev 4, 37, doi:10.1186/1749-8104-4-37 (2009).

47 Griffin R., Binari R. & Perrimon N. Genetic odyssey to generate marked clones in Drosophila mosaics. Proc Natl Acad Sci U S A 111, 4756–4763, doi:10.1073/pnas.1403218111 (2014).

48 Yu H. H., Chen C. H., Shi L., Huang Y. & Lee T. Twin-spot MARCM to reveal the developmental origin and identity of neurons. Nat Neurosci 12, 947–953, doi:10.1038/nn.2345 (2009).

49 Doench J. G. et al. Optimized sgRNA design to maximize activity and minimize off-target effects of CRISPR-Cas9. Nat Biotechnol 34, 184–191, doi:10.1038/nbt.3437 (2016).

50 Haeussler M. et al. Evaluation of off-target and on-target scoring algorithms and integration into the guide RNA selection tool CRISPOR. Genome Biol 17, 148, doi:10.1186/s13059-016-1012-2 (2016).

51 Chari R., Mali P., Moosburner M. & Church G. M. Unraveling CRISPR-Cas9 genome engineering parameters via a library-on-library approach. Nat Methods 12, 823–826, doi:10.1038/nmeth.3473 (2015).

52 Chari R., Yeo N. C., Chavez A. & Church G. M. sgRNA Scorer 2.0: A Species-Independent Model To Predict CRISPR/Cas9 Activity. ACS Synth Biol 6, 902–904, doi:10.1021/acssynbio.6b00343 (2017).

53 Moreno-Mateos M. A. et al. CRISPRscan: designing highly efficient sgRNAs for CRISPR-Cas9 targeting in vivo. Nat Methods 12, 982–988, doi:10.1038/nmeth.3543 (2015).

